# Prediction and Classification of the Uptake and Disposition of Antidepressants and New CNS-Active Drugs in the Human Brain using the ANDROMEDA by Prosilico Software and Brainavailability-Matrix

**DOI:** 10.1101/2022.09.28.509936

**Authors:** Urban Fagerholm, Sven Hellberg, Jonathan Alvarsson, Ola Spjuth

**Affiliations:** Prosilico AB, Lännavägen 7, SE-141 45 Huddinge, Sweden; Department of Pharmaceutical Biosciences and Science for Life Laboratory, Uppsala University, Box 591, SE-751 24 Uppsala, Sweden

## Abstract

**Background:** Passive blood-brain barrier permeability (BBB P_e_), fraction bound to brain tissue (f_b,brain_) and efflux by transport proteins MDR-1 and BCRP are essential determinants for the brain uptake and disposition of drugs.

**Methods:** The main objective of the study was to use the software ANDROMEDA by Prosilico to predict passive BBB P_e_- and f_b,brain_-classes and MDR-1- and BCRP-specificities for various classes of antidepressants and for CNS-active small drugs marketed during 2020 and 2021, and then to position them according to a new 2-dimensional Brainavailability-Matrix (8 passive BBB P_e_ x 4 f_b,brain_ classes, where class 11 has highest and 84 lowest values/brainavailability). Predicted estimates were used, except for cases where measured values were available.

**Results and Conclusion:** Results for 53 drugs show that adequate CNS uptake and disposition are achieved for compounds placed in the zones for low, moderate and high brainavailability, despite efflux. They also show that high brainavailability and efflux are common for CNS-active drugs and that modern CNS-active drugs generally have lower brainavailability than older antidepressive drugs. Furthermore, they demonstrate that ANDROMEDA by Prosilico and the new Brainavailability-Matrix are applicable for prediction, optimization and positioning of CNS uptake and disposition of drugs and drug candidates in man.

## Introduction

Various parameters are used for the estimation, description and prediction of the uptake and disposition of drugs in the brain, for example, fraction bound to brain tissue (f_b,brain_), blood-brain barrier permeability (BBB P_e_), brain-to-plasma concentration ratio (K_p,brain_) and the unbound brain-to-plasma concentration ratio (K_p,uu,brain_).

The uptake into the brain is favored by a high passive BBB P_e_ and absence of efflux by transport proteins such as MDR-1 and BCRP, and the disposition is favored by both a high total BBB P_e_ and a high f_b,brain_.

It is difficult to measure uptake and central nervous system (CNS)-disposition of drugs *in vivo* in man. Instead, data from studies in cells and animals and *in silico* models are used to predict them.

Points of view that K_p,brain_ is a measurement of BBB P_e_ and that the BBB is impermeable, or at least a major obstacle, for compounds with certain characteristics influences the interpretation of the “brainavaialability” (BBB uptake and brain binding) of compounds. K_p,brain_ is also determined by the binding capacity of compounds to blood components and brain tissue, and is, therefore, not a reliable measurement of BBB P_e_. A critical review showed that the BBB is highly permeable, and sufficiently permeable to absorb compounds with molecular weight (MW) of at least 1000 g/mole (inulin with a MW of ~5000 g/mole and octapeptides with MWs exceeding 1000 g/mole are absorbed across the rat BBB) and a polar surface area (PSA) above 120 Å^2^ and compounds with log D<-3.5 (1).

Permeability is a rate, which means that any compound with a permeability will eventually cross cell membranes and cells. The transit times across the BBB for caffeine and propranolol (high P_e_), morphine (moderate P_e_) and sucrose and inulin (low P_e_) have been estimated to ~0.2 s, ~0.3 s, 7 s, 12 min and 50 min, respectively (1).

Our prediction software ANDROMEDA by Prosilico predicts human clinical pharmacokinetics (PK), including BBB P_e_-class (8 classes; a high Q^2^ has previously been shown for our passive P_e_-model (data on file)), f_b,brain_-class (4 classes; assuming that binding to human brain tissue resembles binding to rat brain homogenate; a Q^2^ of 0.75 for log predictions *vs* observations was achieved with our *in silico* methodology) and efflux by BBB transporters MDR-1 and BCRP. The software is based on conformal prediction (CP) methodology, which is a methodology that sits on top of machine learning methods and produce valid levels of confidence (2), unique algorithms and a new human physiologically-based pharmacokinetic (PBPK) model (3). For a more extensive introduction to CP we refer to Alvarsson et al. 2014 (4). Its major domain is for molecules with molecular weight (MW) of 100-700 g/mole, but it has been demonstrated to predict human clinical PK for compounds with MW up to ~1200-1300 g/mole well (5–7).

The main objective of the study was to use ANDROMEDA to predict passive BBB P_e_- and f_b,brain_-classes and MDR-1- and BCRP-specificities for various classes of antidepressants and for CNS-active small drugs marketed during 2020 and 2021, and then to position them according to a 2-dimensional Brainavailability-Matrix (passive BBB P_e_-class *vs* f_b,brain_-class). The use of such a matrix is believed to make interpretations of CNS-uptake and -disposition of drugs and drug candidates simpler.

## Methods

### Selection of CNS-active compounds

34 antidepressants from various classes (including SSRIs, SNRIs, TCAs, and benzodiazepines) and 19 small drugs (including one prodrug) with CNS-activity marketed during 2020 and 2021 were selected. The MW of these compounds are maximally 1117 g/mole (setmelanotide) and 4 of them has a MW above 600 g/mole (atogepant, difelikefalin, lonafarib and setmelanotide).

### Predictions and classifications of CNS uptake and disposition

The prediction system based on CP and a new PBPK-model (ANDROMEDA software by Prosilico for prediction, simulation and optimization of human clinical PK) was applied to predict the main parameters for the study: passive BBB P_e_-class (8 classes; high P_e_ for classes 1-3 and highest P_e_ for class 1, moderate P_e_ for classes 4-6 and low P_e_ for for classes 7 and 8), f_b,brain_-class (4 classes; ≥98, 90-97.99, 50-89.99 and <50 % binding for classes 1, 2, 3 and 4, respectively; model developed based on fractional binding to rat brain homogenate), and MDR-1 and BCRP substrate (efflux) specificity.

Compounds were placed in the 2-dimensional Brainavailability-Matrix (8 passive BBB P_e_ x 4 f_b,brain_ classes), with predefined zones for low, moderate and high brainavailability (class 11 has highest brainavailability and class 84 has lowest brainavailability).

For some antidepressants marketed before 2020, f_b,brain_-estimates and efflux substrate specificities were already available. In such cases, measured estimates were used and presented. Predicted values were produced when laboratory data were missing and for all the new small drugs marketed in 2020 and 2021. Every prediction was a forward-looking prediction where each compound was unknown to the models of the software.

## RESULTS

Results based on measurements (~1/3 for f_b,brain_ and efflux of antidepressants) or predictions are shown in Figure 1 (positioning into the Brainavailability-Matrix) and Table 1. 22 (~42 %) drugs were positioned in the predefined high brainavailability zone (green), out of which 8 belong to class 11 (desipramine, duloxetine, fluoxetine, maprotiline, nefazodone, nortriptyline, sertraline and vortioxetine). All of these with high brainavailability (maybe with the exception of drug #43, sibutramine) are effluxed by MDR-1 and/or BCRP (according to predictions). 30 (~57 %) drugs were placed in the region for moderate (yellow) predefined brainavailability, and only 1 (~2 %) drug (#42, setmelanotide with MW 1117 g/mole was allocated to the low brainavailability zone (orange) (class 82). All of these with low and moderate brainavailability, except for drug #6 (buproprion), were measured or predicted to be effluxed at the BBB.

**Figure 1.**
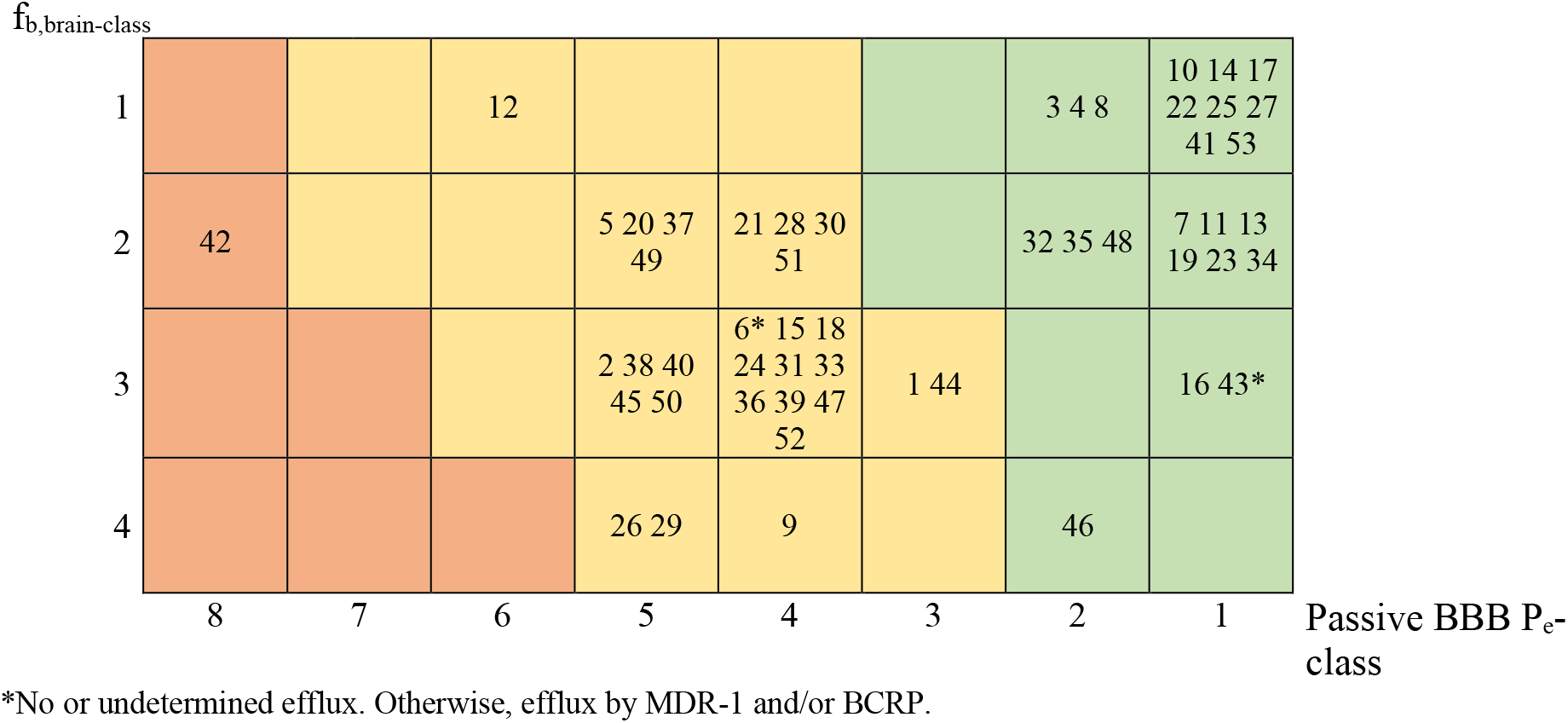
Positioning of CNS uptake and disposition of the 53 drugs (here numbered 1 to 53) into the Brainavailability-Matrix (passive BBB P_e_ *vs* f_b,brain_; green=high; yellow=moderate; orange=low). See Table 1 for drug identities.

**Table 1.** The selected drugs and their measured/predicted BBB P_e_- and f_b,brain_-classes and categories.

| | Drug | Passive<br>BBB $P_e$<br>class | $f_{b,brain}$-<br>binding<br>class | Category |
| --- | --- | --- | --- | --- |
| 1 | Alprazolam | 3 | 3 | Benzodiazepine |
| 2 | Amisulpride | 5 | 3 | Marketed 2020-2021 |
| 3 | Amitriptyline | 2 | 1 | TCA |
| 4 | Amoxapine | 2 | 1 | TCA |
| 5 | Atogepant | 5 | 2 | Marketed 2020-2021 |
| 6 | Bupropion | 4<sup>1</sup> | 3 | Other |
| 7 | Citalopram | 1 | 2 | SSRI/SNRI |
| 8 | Clomipramine | 2 | 1 | TCA |
| 9 | Clonidine | 4 | 4 | Other |
| 10 | Desipramine | 1 | 1 | TCA |
| 11 | Diazepam | 1 | 2 | Benzodiazepine |
| 12 | Difelikefalin | 6 | 1 | Marketed 2020-2021 |
| 13 | Doxepin | 1 | 2 | TCA |
| 14 | Duloxetine | 1 | 1 | SSRI/SNRI |
| 15 | Fexinidazole | 4 | 3 | Marketed 2020-2021 |
| 16 | Flortaucipir F18 | 1 | 3 | Marketed 2020-2021 |
| 17 | Fluoxetine | 1 | 1 | SSRI/SNRI |
| 18 | Fluvoxamine | 4 | 3 | SSRI/SNRI |
| 19 | Imipramine | 1 | 2 | TCA |
| 20 | Lonafarnib | 5 | 2 | Marketed 2020-2021 |
| 21 | Lorazepam | 4 | 2 | Benzodiazepine |
| 22 | Maprotiline | 1 | 1 | TCA |
| 23 | Midazolam | 1 | 2 | Benzodiazepine |
| 24 | Milnacipran | 4 | 3 | SSRI/SNRI |
| 25 | Nefazodone | 1 | 1 | Other |
| 26 | Nifurtimox | 5 | 4 | Marketed 2020-2021 |
| 27 | Nortriptyline | 1 | 1 | TCA |
| 28 | Oliceridine | 4 | 2 | Marketed 2020-2021 |
| 29 | Opicapone | 5 | 4 | Marketed 2020-2021 |
| 30 | Oxazepam | 4 | 2 | Benzodiazepine |
| 31 | Ozanimod | 4 | 3 | Marketed 2020-2021 |
| 32 | Paroxetine | 2 | 2 | SSRI/SNRI |
| 33 | Phenelzine | 4 | 3 | Other |
| 34 | Ponesimod | 1 | 2 | Marketed 2020-2021 |
| 35 | Reboxetine | 2 | 2 | Other |
| 36 | Remimazolam | 4 | 3 | Marketed 2020-2021 |
| 37 | Rimegepant | 5 | 2 | Marketed 2020-2021 |
| 38 | Samidorphan | 5 | 3 | Marketed 2020-2021 |
| 39 | Selumetinib | 4 | 3 | Marketed 2020-2021 |
| 40 | Serdexmethylphenidate<sup>2</sup> | 5 | 3 | Marketed 2020-2021 |
| 41 | Sertraline | 1 | 1 | SSRI/SNRI |
| 42 | Setmelanotide | 8 | 2 | Marketed 2020-2021 |
| 43 | Sibutramine | 1<sup>3</sup> | 3 | SSRI/SNRI |
| 44 | Temazepam | 3 | 3 | Benzodiazepine |
| 45 | Teniloxazine | 5 | 3 | Other |
| 46 | Tramadol | 2 | 4 | SSRI/SNRI |
| 47 | Trazodone | 4 | 3 | OTHER |
| 48 | Triazolam | 2 | 2 | Benzodiazepine |
| 49 | Trilaciclib | 5 | 2 | Marketed 2020-2021 |
| 50 | Venlafaxine | 3 | 3 | SSRI/SNRI |
| 51 | Vilazodone | 4 | 2 | Other |
| 52 | Viloxazine | 4 | 3 | Marketed 2020-2021 |
| 53 | Vortioxetine | 1 | 1 | Other |

<sup>1</sup>No efflux; <sup>2</sup>Prodrug of ADHD-medicine dexamethylphenidate; <sup>3</sup>Undetermined efflux.

Only 2 of the new CNS-active drugs on the market were predicted to have high BBB P_e_. Most of them belong to class 4-5 BBB P_e_. 24 of the older antidepressants were predicted to have high BBB P_e_. The median BBB P_e_ and f_b,brain_ classes are 2 and 2 for antidepressants and 5 and 3 for new CNS-active drugs, respectively.

Figure 2 shows (as an example) predicted molecular regions of ponesimod (brainavailability class 12) contributing to decreasing and increasing the passive BBB P_e_ and f_b,brain_.

**Figure 2.**
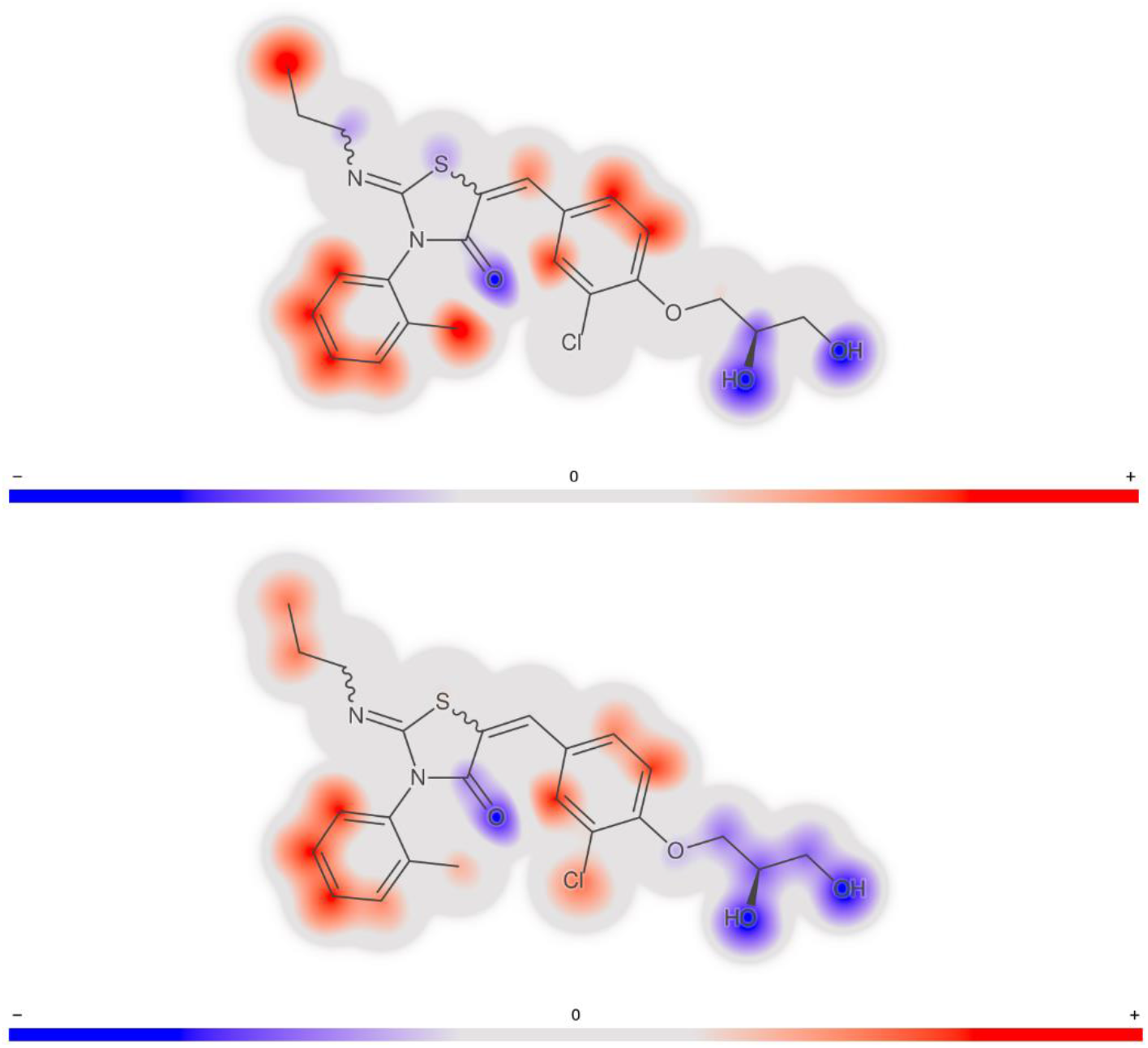
Molecular structure and signatures for ponesimod (brainavailability class 12). Molecular regions contributing to decreasing (blue) and increasing (red) the passive BBB (upper) P_e_ and (lower) f_b,brain_ are shown.

## Discussion & Conclusion

Results show that adequate CNS uptake and disposition were achieved for compounds placed in the zones for low, moderate and high brainavailability, despite efflux by MDR-1 and/or BCRP. CNS-activity is not only dependent on BBB P_e_, but also on the potency, dose and systemic exposure, and this might explain why compounds with low brainavailability can produce sufficient CNS-effects. Alternatively, the BBB is sufficiently Permeable for effluxed low P_e_-compounds with a MW of ~1000 g/mole. Results also show that high brainavailability and efflux are common for CNS-active drugs and that modern CNS-active drugs generally have lower brainavailability than older antidepressive drugs. Furthermore, they demonstrate that ANDROMEDA by Prosilico and the new Brainavailability-Matrix are applicable for prediction, optimization and positioning of CNS uptake and disposition of drugs and drug candidates in man.

